# The transcriptome of *Balamuthia mandrillaris* trophozoites for structure-based drug design

**DOI:** 10.1101/2020.06.29.178905

**Authors:** Isabelle Q. Phan, Christopher A. Rice, Justin Craig, Rooksana E. Noorai, Jacquelyn McDonald, Sandhya Subramanian, Logan Tillery, Lynn K. Barrett, Vijay Shankar, James C. Morris, Wesley C. Van Voorhis, Dennis E. Kyle, Peter J. Myler

**Affiliations:** Seattle Structural Genomics Center for Infectious Disease (SSGCID), Seattle, Washington, USA; Center for Global Infectious Disease Research, Seattle Children’s Research Institute, Seattle, Washington, USA; Center for Tropical and Emerging Global Diseases, University of Georgia, Athens, Georgia, USA; Center for Emerging and Re-emerging Infectious Diseases (CERID), Division of Allergy and Infectious Diseases, Department of Medicine, University of Washington, Seattle, Washington, USA; Clemson University Genomics and Bioinformatics Facility, Clemson University, Clemson, South Carolina, USA; Center for Human Genetics, Clemson University, Greenwood, South Carolina, USA; Eukaryotic Pathogens Innovation Center, Department of Genetics and Biochemistry, Clemson University, Clemson, South Carolina, USA; Department of Microbiology, University of Washington, Seattle, Washington, USA; Department of Global Health, University of Washington, Seattle, Washington, USA; Department of Pediatrics, University of Washington, Seattle, Washington, USA

## Abstract

*Balamuthia mandrillaris*, a pathogenic free-living amoeba (FLA), causes cutaneous skin lesions as well as the brain-eating disease: *Balamuthia* granulomatous amoebic encephalitis (GAE). These diseases, and diseases caused by other pathogenic FLA, *Naegleria fowleri* or *Acanthamoeba* species, are minimally studied from a drug discovery perspective; few targets have been validated or characterized at the molecular level, and little is known about the biochemical pathways necessary for parasite survival. Chemotherapies for CNS disease caused by *B. mandrillaris* require vast improvement. Current therapeutics are limited to a small number of drugs that were previously discovered in the last century through *in vitro* testing or identified after use in the small pool of surviving reports.

Using our recently published methodology to identify potentially useful therapeutics, we screened a collection of 85 compounds that have previously been reported to have antiparasitic activity. We identified 59 compounds that impacted growth at concentrations below 220 μM. Since there is no fully annotated genome or proteome, we used RNA-Seq to reconstruct the transcriptome of *B. mandrillaris* and locate the coding sequences of the specific genes potentially targeted by the compounds identified to inhibit trophozoite growth. We determined the sequence of 17 of these target genes and obtained expression clones for 15 that we validated by direct sequencing.

## Introduction

*Balamuthia mandrillaris* is a ubiquitous soil-dwelling amoeba that is the causative agent of GAE^1–4^. In common with the other two major pathogenic free-living amoebas (FLAs), *Naegleria fowleri* and *Acanthamoeba castellanii*, *B. mandrillaris* infections, though uncommon, have > 90% case fatality rate^5^. In the United States, 109 *Balamuthia* cases in both immunocompetent and immunocompromised individuals have been reported with at least twice that many worldwide, but these rates of GAE are likely to be underestimated due to historically poor diagnosis^6^. Nonetheless, awareness of potential cases has been on the rise^7,8^. In addition to diagnostic awareness, increasing rates of amoebic infections in northern regions of the United States could also be early indicators that recent emergence of these diseases might be associated with global warming^6^. In contrast to *Naegleria* infections that present and progress extremely rapidly after exposure, *Balamuthia* incubation times might be as long as several months and disease progression more subacute or chronic, increasing the opportunity for therapeutic intervention^9^. However, treatment options remain very limited, leading to poor outcomes even with the correct diagnosis.

The recent development of an inexpensive and easily prepared media, as well as increasing interest in *B. mandrillaris* as a public health concern, has facilitated the development of robust high-throughput drug screening methods. Where low throughput methods restricted screening to ~10-20 drugs at a time, the new high-throughput methods allow rapid screening of hundreds to thousands of drugs simultaneously and allows direct comparisons of activity. In this study, we identify FDA approved drugs that could potentially be repurposed for therapy alone or added into a combination drug cocktail against *B. mandrillaris*. These drugs can also be used as leads for further structure-based drug discovery (SBDD) exploration and *in vivo* efficacy studies.

Structure based drug discovery (SBDD) was originally devised in the mid 1980’s^10^. With advances in methodology for protein structure elucidation, less expensive and faster computer processing, and improved access to prediction software, the timeline of solving target structures and developing specific and selective drugs have significantly shortened^10^. SBDD has been discussed by several authors throughout the amoeba literature as an attractive method of designing selective enzymatic inhibitors that would specifically target the parasite over the human host^11^. Given how frequently this strategy is discussed, it is surprising that only a small number of laboratories have actually tested this methodology in practice against pathogenic FLA. Sterol biosynthesis has been the most attractive target since parasites utilize ergosterol over cholesterol for making cell plasma membranes with distinct host biosynthetic differences that could be selectively targeted^12–14^. Glucose metabolism is essential for parasitic cell viability. Milanes *et al*., recently targeted glucokinase in *Naegleria fowleri*, and described NfGlck specific inhibitors with minimal activity against human glucokinase in recombinant enzymatic functional studies^15^. Other studies have looked at targeting histidine or shikimate essential amino acid biosynthetic pathways in *Acanthamoeba* species, which the hosts cannot synthesize *de novo,* as parasite specific targets for drug intervention^16,17^.

The development of new compounds against *B. mandrillaris* in particular has been hampered by the paucity of genomic information. Though draft genomes have been published, no structural and functional annotation is currently available^18,19^. This information is essential for the design of new drugs by SBDD, as that methodology requires information about the molecular structure of the target protein. Once the protein coding sequences are annotated on the genome, rapid selection of multiple drug targets can be performed, for example by homology searches with known drug targets, thus paving the way for combinational therapy, a broadly established strategy to minimize the risk of drug resistance. This study presents the first comprehensive proteome of *B. mandrillaris* reconstructed from RNA sequencing of logarithmic growing trophozoites, the infective form of the amoeba. Potential drug targets identified through phenotypic screening were selected specifically from the trophozoite transcriptome and PCR amplified. The clones were further validated by direct sequencing, providing the first step for recombinant expression and crystallization by the Seattle Structural Genomics Center for Infectious Disease (SSGCID) high-throughput gene-to-structure pipeline^20^.

## Results and discussion

### Phenotypic screens

We performed a drug susceptibility screen of 85 compounds, previously identified as antiparasitic, against the trophozoite stage of *B. mandrillaris*, and discovered that 59 of these compounds had 50% inhibitory concentration (IC_50_) efficacy at ≤ 220 μM concentration (**Table 1**). Two compounds (dequalinium choride and alexidine) possessed nanomolar potency. We found that many of the current drugs used within the treatment regimen for *Balamuthia* GAE appeared to be only moderately to slightly efficacious, with IC_50_ activity ranging from 18.35 μM (pentamidine) to > 163.25 μM (fluconazole). Here we should also note that miltefosine, the newest drug addition to the amoebae chemotherapy cocktail, was inactive at the final screening concentration of > 122.68 μM.

**Table 1.**
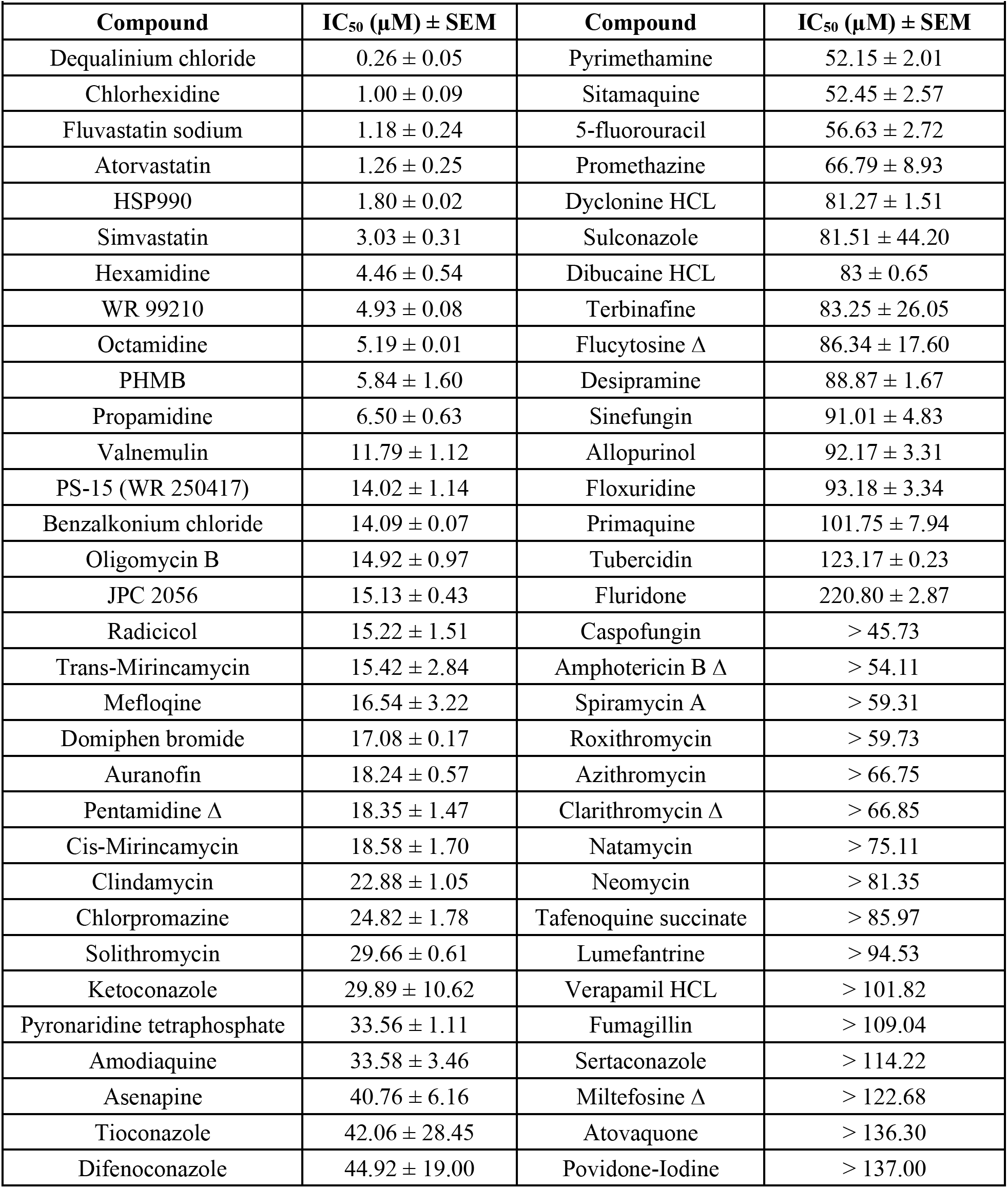
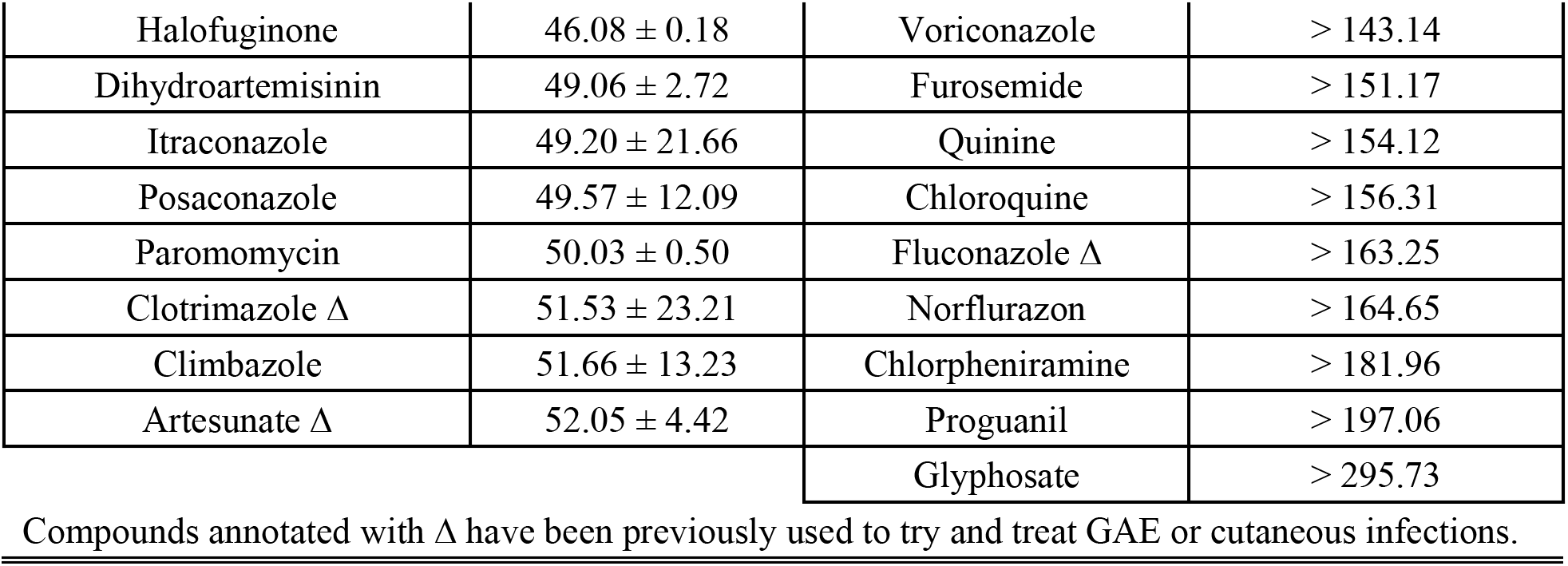
Phenotypic analysis of 85 compounds against logarithmic trophozoites *in vitro*. Compounds annotated with Δ were previously used within known patients’ treatment regimens for *Balamuthia* GAE or cutaneous *Balamuthia* infections. The susceptibility is ranked in order of highly potent (left hand side column) to minimal potency (right hand side column) and the inhibitory concentration that causes 50% ATP depletion (death) is listed (IC_50_). All compounds were initially screened from 50 μg/ml and converted to molarity for standardized testing.

| Compound | IC<sub>50</sub> (μM) ± SEM | Compound | IC<sub>50</sub> (μM) ± SEM |
| --- | --- | --- | --- |
| Dequalinium chloride | 0.26 ± 0.05 | Pyrimethamine | 52.15 ± 2.01 |
| Chlorhexidine | 1.00 ± 0.09 | Sitamaquine | 52.45 ± 2.57 |
| Fluvastatin sodium | 1.18 ± 0.24 | 5-fluorouracil | 56.63 ± 2.72 |
| Atorvastatin | 1.26 ± 0.25 | Promethazine | 66.79 ± 8.93 |
| HSP990 | 1.80 ± 0.02 | Dyclonine HCL | 81.27 ± 1.51 |
| Simvastatin | 3.03 ± 0.31 | Sulconazole | 81.51 ± 44.20 |
| Hexamidine | 4.46 ± 0.54 | Dibucaine HCL | 83 ± 0.65 |
| WR 99210 | 4.93 ± 0.08 | Terbinafine | 83.25 ± 26.05 |
| Octamidine | 5.19 ± 0.01 | Flucytosine $\Delta$ | 86.34 ± 17.60 |
| PHMB | 5.84 ± 1.60 | Desipramine | 88.87 ± 1.67 |
| Propamidine | 6.50 ± 0.63 | Sinefungin | 91.01 ± 4.83 |
| Valnemulin | 11.79 ± 1.12 | Allopurinol | 92.17 ± 3.31 |
| PS-15 (WR 250417) | 14.02 ± 1.14 | Floxuridine | 93.18 ± 3.34 |
| Benzalkonium chloride | 14.09 ± 0.07 | Primaquine | 101.75 ± 7.94 |
| Oligomycin B | 14.92 ± 0.97 | Tubercidin | 123.17 ± 0.23 |
| JPC 2056 | 15.13 ± 0.43 | Fluridone | 220.80 ± 2.87 |
| Radicicol | 15.22 ± 1.51 | Caspofungin | > 45.73 |
| Trans-Mirincamycin | 15.42 ± 2.84 | Amphotericin B $\Delta$ | > 54.11 |
| Mefloquine | 16.54 ± 3.22 | Spiramycin A | > 59.31 |
| Domiphen bromide | 17.08 ± 0.17 | Roxithromycin | > 59.73 |
| Auranofin | 18.24 ± 0.57 | Azithromycin | > 66.75 |
| Pentamidine $\Delta$ | 18.35 ± 1.47 | Clarithromycin $\Delta$ | > 66.85 |
| Cis-Mirincamycin | 18.58 ± 1.70 | Natamycin | > 75.11 |
| Clindamycin | 22.88 ± 1.05 | Neomycin | > 81.35 |
| Chlorpromazine | 24.82 ± 1.78 | Tafenoquine succinate | > 85.97 |
| Solithromycin | 29.66 ± 0.61 | Lumefantrine | > 94.53 |
| Ketoconazole | 29.89 ± 10.62 | Verapamil HCL | > 101.82 |
| Pyronaridine tetraphosphate | 33.56 ± 1.11 | Fumagillin | > 109.04 |
| Amodiaquine | 33.58 ± 3.46 | Sertaconazole | > 114.22 |
| Asenapine | 40.76 ± 6.16 | Miltefosine $\Delta$ | > 122.68 |
| Tioconazole | 42.06 ± 28.45 | Atovaquone | > 136.30 |
| Difenoconazole | 44.92 ± 19.00 | Povidone-Iodine | > 137.00 |
| Halofuginone | 46.08 ± 0.18 | Voriconazole | > 143.14 |
| Dihydroartemisinin | 49.06 ± 2.72 | Furosemide | > 151.17 |
| Itraconazole | 49.20 ± 21.66 | Quinine | > 154.12 |
| Posaconazole | 49.57 ± 12.09 | Chloroquine | > 156.31 |
| Paromomycin | 50.03 ± 0.50 | Fluconazole Δ | > 163.25 |
| Clotrimazole Δ | 51.53 ± 23.21 | Norflurazon | > 164.65 |
| Climbazole | 51.66 ± 13.23 | Chlorpheniramine | > 181.96 |
| Artesunate Δ | 52.05 ± 4.42 | Proguanil | > 197.06 |
|  |  | Glyphosate | > 295.73 |

Compounds annotated with Δ have been previously used to try and treat GAE or cutaneous infections.

Based on previously determined *in vitro* activity and the few surviving cases of *Balamuthia* GAE infections, the Centers for Disease Control and Prevention (CDC) recommends that the drug cocktail regimen for treating disease include a combination of pentamidine, sulfadiazine, flucytosine, fluconazole, azithromycin or clarithromycin, and miltefosine^6^. We thus proceeded to test these compounds, starting with the macrolides. Our screening results consistently indicate that the compounds belonging to the macrolide drug class (azithromycin, clarithromycin, roxithromycin, and spiramycin) are inactive, in agreement with previous results^21^. Interestingly, solithromycin, a known ketolide antibiotic against macrolide-resistant Streptococcal species^22^, appeared to show moderate activity against *B. mandrillaris* (29.66 μm). We found that other macrolides such as amphotericin B and natamycin, examples of polyene antimycotics, a subgroup of macrolides, that are generally used for targeting ergosterol within fungal cell membranes^23^, were also inactive against *B. mandrillaris*. We then tested the azole compounds. CDC studies reported that fluconazole was inactive at concentrations lower than 10 μg/ml^24^. We were able to confirm the fluconazole result; however, other antifungal azoles (ketoconazole, tioconazole, difenoconazole, itraconazole, posaconazole, clotrimazole, climbazole, and sulconazole) displayed better, though still moderate, activity (29.89-81.51 μm). We tested flucytosine and miltefosine and both displayed moderate to poor activity against *B. mandrillaris* with an IC_50_ of 86.34 and > 122 μm, respectively. Flucytosine was previously described at 10 μg/ml (77 μm) to inhibit 61% *Balamuthia* cytopathogenicity, these and our results suggest an equipotency agreement^25^. Although miltefosine has been reported to have moderate activity at concentrations of 63-100 μm^25,26^, another study^27^ as well as ours suggest miltefosine activity may be less potent with a higher IC_50_ of >122 μm. As for pentamidine, our results are consistent with prior studies that obtained comparable IC_50_ values between 9 and 29 μm^24,28,29^. Thus, of the combination drug cocktail recommended by the CDC, only pentamidine and flucytosine appears to have *in vitro* activity against *B. mandrillaris,* though we were unable to test sulfadiazine. It is possible that the recommended drug therapy is active in combination, and not when tested in isolation as in this study. It is also possible that the drugs are biologically activated *in vivo* or are only active against an *in vivo* form that we have not assayed. We therefore cannot rule out activity for the recommended drugs based solely on our *in vitro* sensitivity screens.

We previously identified statins, which target 3-hydroxy-3-methylglutaryl-coenzyme A reductase A (HMG-CoA), as active compounds against *B. mandrillaris*^29^. As part of this study, we tested three additional statins: fluvastatin, atorvastatin and simvastatin and observed that they were active against *B. mandrillaris* at 1.18 to 3.03 μM concentration. Simvastatin and fluvastatin in particular have been shown to have better brain penetration compared to other statins^30^, indicating potential for off-label drug repurposing.

From a drug-repurposing standpoint, the compounds described in this study yielded a plethora of potentially useful drugs that act on *B. mandrillaris*. However, none of the active compounds in our screens has known mechanisms of action in *Balamuthia* where the specific protein target has been identified. To increase the pool of potential protein targets, we incorporated results from our previous drug discovery studies. These include screening the MMV Malaria and Pathogen boxes that identified 11 compounds with equipotency to nitroxolone (8-Hydroxy-5-nitroquinoline) IC_50_ of 2.84 μM^25^, indicating these molecules could be used as an initial starting point for medicinal chemistry structure activity relationship (SAR) studies. More recently, we identified 63 compounds through screening the Calibr ReFRAME drug repurposing library, with activities ranging from 40 nM to 4.8 μM^29^. Chemical inference from our previous drug screening efforts and the results from this study identified a total of 52 potential protein targets in *B. mandrillaris*. These were reduced to 25 after excluding kinases. These 25 protein targets and an additional six community targets of interest were submitted for structural determination to the Seattle Structural Genomics Center for Infectious Diseases (SSGCID) gene-to-structure pipeline.

The first step of the SSGCID pipeline involves cloning of the *B. mandrillaris* sequences encoding the protein targets identified in the screens. However, lack of annotation of the *Balamuthia* genome hampered these efforts: although we were able to locate some sequences using BLAST searches of the *Balamuthia* genome using the human and *Acanthamoeba* homologues, PCR amplification from *B. mandrillaris* cDNA (or gDNA) was not successful. Therefore, transcriptome sequencing of *Balamuthia* was performed.

### Transcriptome sequencing, assembly and functional annotation

We reconstructed a haploid version of *B. mandrillaris* proteome and estimated that our set of 14.5K proteins was 90% complete according to standard benchmark. Comparison with other species showed highest sequence similarity (averaging 45%) to *A. castellanii* strain Nef. We obtained functional annotation for over 80% of the proteins, and this information will serve as a reference for future functional studies as well as a source for target selection.

The sequencing of RNA isolated from an axenic laboratory culture of *B.mandrillaris* trophozoites yielded 30,473,902 paired-end reads (2 × 75 bp). To build the proteome and compensate for the relatively short RNA-seq reads, we first performed both *de-novo* and genome-based assemblies and then predicted protein coding sequences (CDSs) with EvidentialGene (EviGene)^31^. In this hybrid approach, we combined two *de-novo* assemblies, obtained from different assembly packages and multiple k-mers, and a genome-based assembly based on the existing *B. mandrillaris* genome LFUI01 (**Figure 1**)^19^.

**Figure 1:**
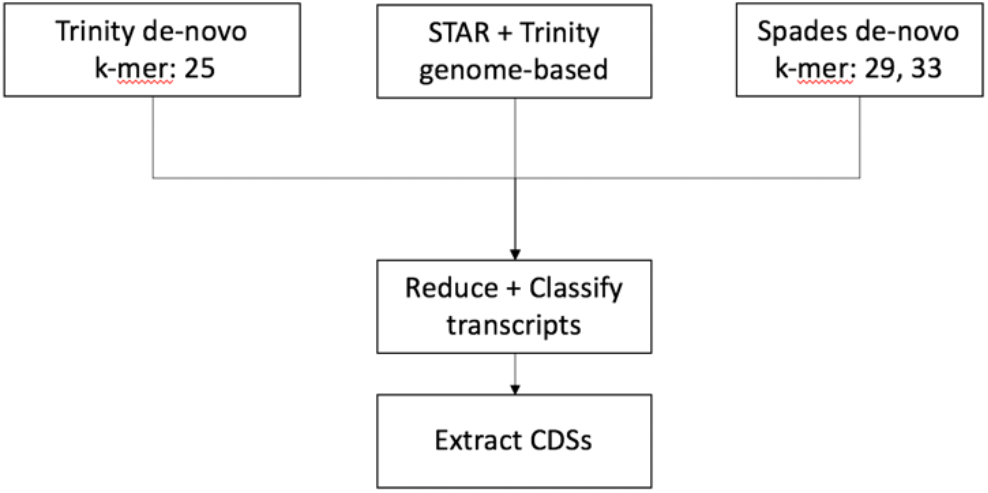
Overview of the main steps for predicting the *B. mandrillaris* proteome from RNA-seq reads using a hybrid approach. *De-novo* and genome-based assemblies are combined and processed with EviGene to reduce transcript redundancy and classify transcripts as encoding complete or incomplete CDSs (5’ and/or 3’ truncated). CDSs are extracted, translated and annotated as “main” or alternate.

This approach yielded a total of 37,252 transcripts, of which 20,005 contained complete CDSs. All CDSs were extracted, translated and resulting protein sequences annotated as either ‘main’ or alternate. The average length of the top 1,000 longest complete proteins was 1,552 ± 387 amino acids, a number indicative of assembly quality that is roughly comparable to the corresponding value derived from the re-annotated AmoebaDBv44 *A. castellanii* proteome (1,688 ± 539)^32,33^. EviGene classified a total of 14,492 sequences as “main” and we use this number as a proxy for estimating the size of the trophozoite haploid proteome. Although less that two-third of the EviGene “main” sequences were complete, they represented 90% of complete eukaryotic Benchmarking Universal Single-Copy Orthologs (BUSCOs), a standard measure to quantify relative accuracy and completeness (**Table 2**)^34^. Table 2 shows comparable levels of completeness for the transcriptome and the unannotated genome (LFUI01), but single copy BUSCO numbers indicate that the EviGene “main” proteins likely represent the haploid proteome.

**Table 2.** Quality and completeness assessment of the assembled transcriptome, the EviGene “main” proteins and the draft genome relative to the dataset for eukaryotes eukaryota_odb9. Input sequences for retrieving BUSCOs are proteins for the proteomes and scaffolds for the genomes. Note: the reduction of duplicates in the set of ‘main’ proteins compared to the full assembly to under 1%, at a marginal cost of 4% loss of complete BUSCOs.

|  | Assembled transcriptome |  | EviGene "main" proteins |  | Genome (LFUI01) |  |
| --- | --- | --- | --- | --- | --- | --- |
| # input sequence | 37,252 |  | 14,492 |  | 1,605 |  |
| Complete BUSCOs (C) | 286 | 94% | 272 | 90% | 271 | 89% |
| single-copy (S) | 92 | 30% | 270 | 89% | 166 | 55% |
| duplicated (D) | 194 | 64% | 2 | 1% | 105 | 35% |
| Fragmented BUSCOs (F) | 4 | 1% | 6 | 2% | 10 | 3% |
| Missing BUSCOs (M) | 13 | 4% | 25 | 8% | 22 | 7% |

As expected, comparison to other species in the UniProt database with AAI-profiler indicates that the closest sequenced proteome is from *A. castellanii* strain Neff, although with only 27% of matched fraction, the two proteomes appear to be distantly related (**Figure 2A**). Note that the AAI-profiler does partial sampling as it relies on SANSparallel, a fast homology search that is as sensitive as BLAST above ca. 50% sequence identity^35^. A BLASTP search of EviGene ‘main’ proteins against *A. castellanii* returned hits for 65% of the *Balamuthia* sequences with an average identity of 44%. Only 38% of hits had over 50% identity, indicating that just about a third of the two proteomes overlap. Indeed, orthologous cluster analysis with *Dictyostelium discoideum* as the outgroup shows that 37% of the *Balamuthia* proteins cluster with *Acanthamoeba,* of which 21% are shared between the three species (**Figure 2B**). This result and the high proportion of singletons (48%) highlights the divergence of *Balamuthia* from *Acanthamoeba*. To place the Balamuthia proteome in an evolutionary context, a neighbour-joining tree was constructed from an alignment-free comparison of complete proteomes from selected Amoebas, with the non-Amoebozoa *Naegleria* as outgroup (**Figure 2C**). As detected by AAI-profiler, the Discosea genera *Balamuthia* and *Acanthamoeba* are in a separate group from the Variosea genus *Planoprotostelium* and the Eumycetozoa Dictyostelids *Cavenderia*, *Polysphondylium*, *Tieghemostelium* and *Dictyostelium*. The Evosea genus *Entamoeba* is in a separate branch from the other Amoebozoa in the tree.

**Figure 2A:**
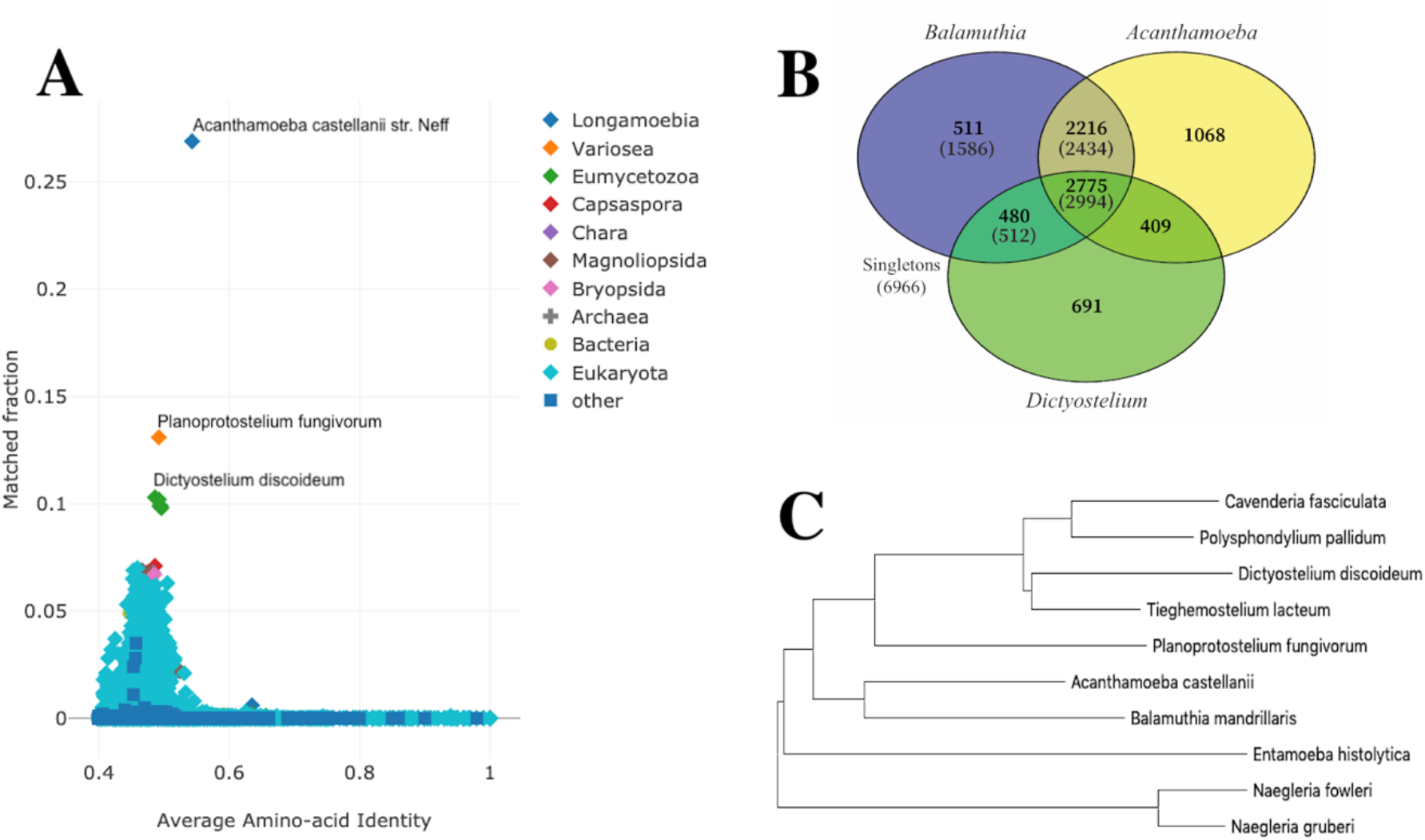
**AAI-profiler scatterplot** of UniProt species with greater than 40% average amino-acid identity to the *Balamuthia* ‘main’ proteins. The species name of the top three proteomes with the largest fraction of matches to *Balamuthia* are indicated. **Figure 2B: Venn diagram** showing the overlap between orthologous cluster groups in the proteomes of *B. mandrillaris*, *A. castellanii* and *D. discoideum*. Total numbers of *B. mandrillaris* proteins in each group are in parenthesis. **Figure 2C: Neighbour-joining tree.** The closest Amoebozoa species was *Acanthamoeba* detected by AAI-profiler with two *Naegleria* and one *Entamoeba* species as outgroups, based on alignment-free comparisons of complete proteomes.

To characterize the proteome further and expand the pool of potential targets, we conducted preliminary functional annotations of the haploid proteome dataset. Functional annotation of the EviGene “main” protein sequences with PANNZER2, one of the top-10 rapid methods in the CAFA2 NK-full benchmark, provided 23% of the sequences with a description and 63% with a lower level GO molecular function term (40% describing a specific activity). A plot of high-level GO terms compared with those obtained for *A. castellanii* and *D. discoideum*, one of the most thoroughly annotated amoebas in UniProt, shows a similar profile for the three species, with differences limited to smaller gene families representing less than 1% of the genes (**Figure 3**). With the caveat that the absence of a gene could be due to assembly errors, *Balamuthia* as well as *Acanthamoeba* appear to be comparatively depleted of genes associated with ubiquitin-mediated protein tagging. The biological implications are unclear. In contrast, the apparent depletion of adhesion genes in *Acanthamoeba* is surprising given the central role of host cell adhesion in *A. castellanii* pathogenesis^36^, even though the absence of Sib cell-adhesion proteins has been reported previously^32^. In terms of potential targets for SBDD, the *Balamuthia* GO annotations classified 858 (5%) of proteins as having kinase activity, of which about half (424) were classified as protein kinases. Our phenotypic screens identified potential kinase targets, but further analysis is needed to determine their specific sequence and target the kinome with accuracy^37^.

**Figure 3:**
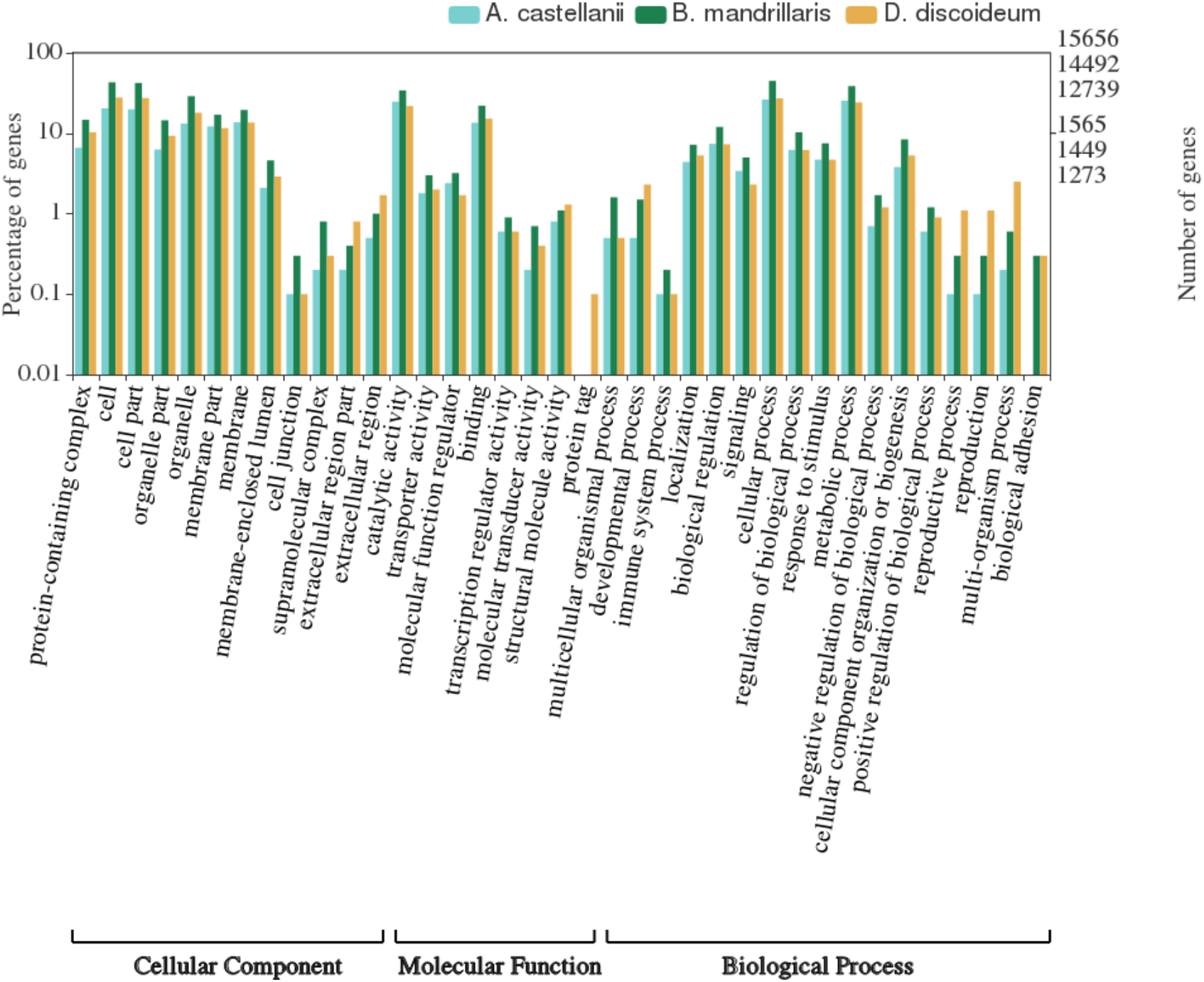
Level 2 top GO annotations. *B. mandrillaris* proteins (dark green) vs *A. castellanii* (cyan) and *D. discoideum* (orange) as percentage of genes and total number of genes on a log(10) scale, significant relationships p-value < 0.05.

### Target identification and validation

For this study, protein kinases from the phenotypic screens were left out, leaving a total of 25 potential targets, to which we added 6 known drug targets requested from the amoeba community. From this list of 31 targets, 19 could be assigned to specific human protein sequences. A total of 14 *Balamuthia* sequences for 13 targets (there are 2 copies of topoisomerase II) were identified from a BLASTP search with the human sequences: 12 from the screens, and 2 known drug targets. Average pairwise identity was 49% with 77% coverage. Another three of the known drug targets were not detected by BLASTP searches of the *Balamuthia* proteome using the human sequences, therefore *Acanthamoeba* sequences were used instead. This yielded a total of 17 *Balamuthia* sequences that were entered into the SSGCID gene-to-structure pipeline. Truncations around putative catalytic domains were designed for 9 of the 17 sequences to increase crystallization likelihood, leading to 23 constructs as cloning candidates. PCR amplification produced clones for 18 constructs. Direct sequencing was successful for 15 of these and sequence comparison with the “main” proteins from the EviGene assembly showed excellent matches with over 99% average amino acid identity, corresponding to 2 amino acid variations on average per sequence, and 100% coverage for all, but the two largest proteins (84% coverage and 100% identity for the 1,068 amino-acid long Exportin-1, 81% coverage and 99% identity for the 784 residue primase and C-term domains of topoisomerase II).

The identity between the 15 validated protein sequences and their closest *A. castellanii* homologue ranged between 56% to 88%, with 3 notable exceptions; exportin-1 (21% identity), lanosterol 14-alpha demethylase (CYP51A) (28% identity) and glucokinase (51% identity). In the case of exportin-1, a multiple sequence alignment indicated that the *Balamuthia* protein was over 50% identical to the *Naegleria fowleri* and *Planoprotostelium fungivorum* proteins, suggesting potential mis-assembly in the *A. castellanii* genome. Similarly, the *Acanthamoeba* glucokinase sequence appears to have a large deletion of over 30 residues compared to the *Balamuthia* and *Naegleria* sequences. This region corresponds to a double-stranded beta-sheet that lies in the glucose binding pocket in the *Naegleria* structure, and we would expect it to be conserved in *Acanthamoeba* (**Figure 4**)^15^. As for CYP51A, pairwise identity between the closest *Balamuthia*, *Acanthamoeba* and *Naegleria* sequences ranged between 20% and 24%, suggesting a large divergence in this protein family. Hence, with the exception of CYP51A, the validated targets from *Balamuthia* are likely to share similar binding pockets in *A. castellanii*, as they lie within the overall 55% sequence identity threshold that was shown to be associated with a conserved active site in bacteria^38^. However, the same threshold might not apply to Eukaryotic enzymes.

**Figure 4:**
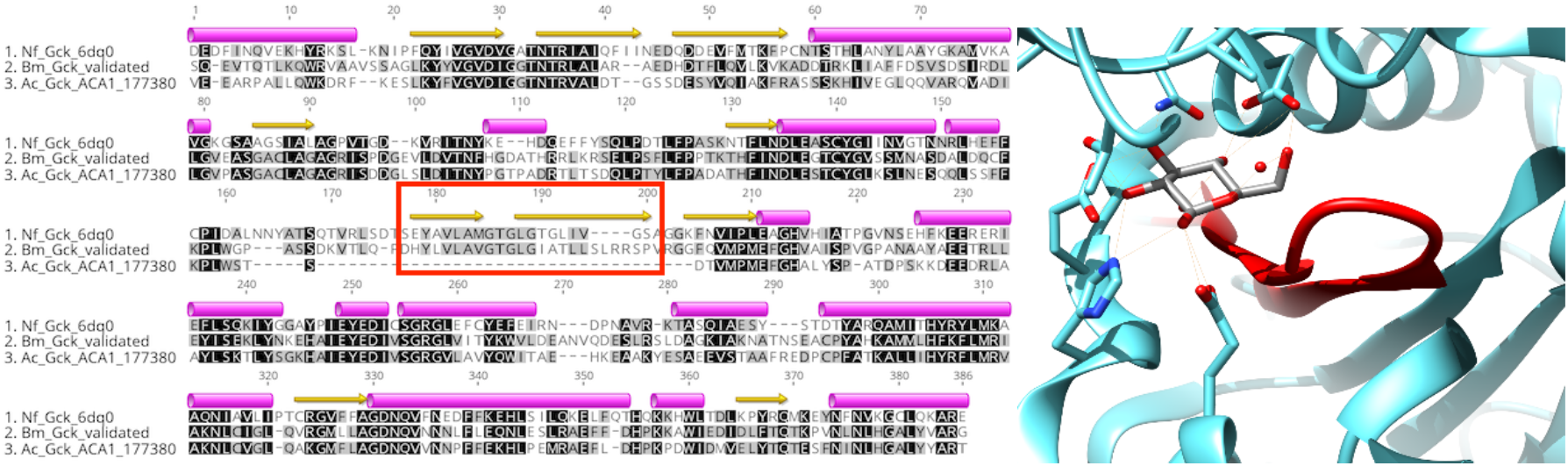
Sequence conservation of glucokinase in 3 pathogenic amoebas. Potential misassembly of the *A. castellanii* glucokinase (AmoebaDB ACA1_177380) highlighted on a multiple sequence alignment with the *B. mandrillaris* validated sequence (this study) and *N. fowleri* crystal structure (PDB: 6DA015; helical regions are annotated as pink tubes and beta-sheets as yellow arrows. The alignment was obtained with T-Coffee-Expresso39. The double-stranded beta-sheet missing in *A. castellanii* glucokinase is colored in red on the active site of the *B. mandrillaris* structure (PDB: 6VZZ)

Of the 13 targets that were also found in human, five shared over 55% sequence identity overall to their human counterpart and might potentially have similar active sites: S-adenosyl-homocysteinase (SAHH), 3-hydroxy-3-methylglutaryl-coenzyme A reductase (HMGR), heat-shock protein 90-alpha (HSP90ɑ), histone deacetylase 1 (HDAC1) and exportin-1 (XPO1) (**Table 3**). As a consequence, SBDD for these targets will likely require exploration of potential alternate binding sites that are specific to the *Balamuthia* protein. We would expect selectivity to be more readily achievable for the other targets, with the topoisomerase ATPases as borderline cases. One promising example of a *Balamuthia* target that can be selectively targeted is the GARTFase domain of trifunctional purine biosynthetic protein adenosine-3 (GART). *Balamuthia* GARTFase has a low sequence identity to the human enzyme (37%) and has a different domain arrangement than in human GART. Whereas GARTFase is the C-terminal domain of human GART, it is the middle domain in *Balamuthia* (**Figure 5**). This domain arrangement, confirmed by direct sequencing and conserved in *Acanthamoeba*, leads us to postulate that targeting double domains in GART may offer a promising avenue to develop drugs against those pathogenic amoebas.

**Table 3.** Sequence similarity (BLASTP) between *Balamuthia* validated sequences and UniProt identifiers of closest human counterpart. (*) indicate lower than expected coverage due to incomplete sequencing of the clones; (**) indicate additional targets selected independently of the Calibr screens. Note that homology to human Glucokinase was too low to be detected with BLASTP at the chosen E-value (1e-3).

| Balamuthia target | Pairwise identity | Target coverage | Closest human protein |
| --- | --- | --- | --- |
| S-adenosyl-L-homocysteine hydrolase | 58% | 98% | sp\|P23526\|SAHH_HUMAN |
| Histone deacetylase 1 | 73% | 83% | sp\|Q13547\|HDAC1_HUMAN |
| Lanosterol 14-alpha demethylase (CYP51A)** | 26% | 96% | sp\|Q16850\|CP51A_HUMAN |
| Methionyl-tRNA synthetase (methionine tRNA ligase) (MetRS)** | 54% | 79% | sp\|P56192\|SYMC_HUMAN |
| Heat shock protein HSP90-alpha | 69% | 100% | sp\|P07900\|HS90A_HUMAN |
| Calcium ATPase, haloacid dehydrogenase (HAD) domain | 43% | 100% | tr\|A0A0A0MSP0\|ATP2C2_HUMAN |
| 3-hydroxy-3-methylglutaryl-CoA reductase (HMG-CoA reductase) (HMGR) | 62% | 97% | sp\|P04035\|HMDH_HUMAN |
| Glucokinase** | - | - | none |
| DNA topoisomerase II copy 1, ATPase and transducer domains | 55% | 97% | sp\|Q02880\|TOP2B_HUMAN |
| DNA topoisomerase II copy 1, toprim and C-term domains | 49% | 99% | sp\|P11388\|TOP2A_HUMAN |
| DNA topoisomerase II copy 2, ATPase and transducer domains | 52% | 97% | sp\|P11388\|TOP2A_HUMAN |
| DNA topoisomerase II copy 2, toprim and C-term domains | 39% | 80%* | sp\|Q02880\|TOP2B_HUMAN |
| Exportin-1 (CRM1, XPO1) | 57% | 83%* | sp\|O14980\|XPO1_HUMAN |
| Xylose isomerase (xylA) | - | - | none |
| Trifunctional purine biosynthetic protein adenosine-3 (GART), GARTFase domain | 37% | 82% | sp\|P22102\|PUR2_HUMAN |

**Figure 5:**
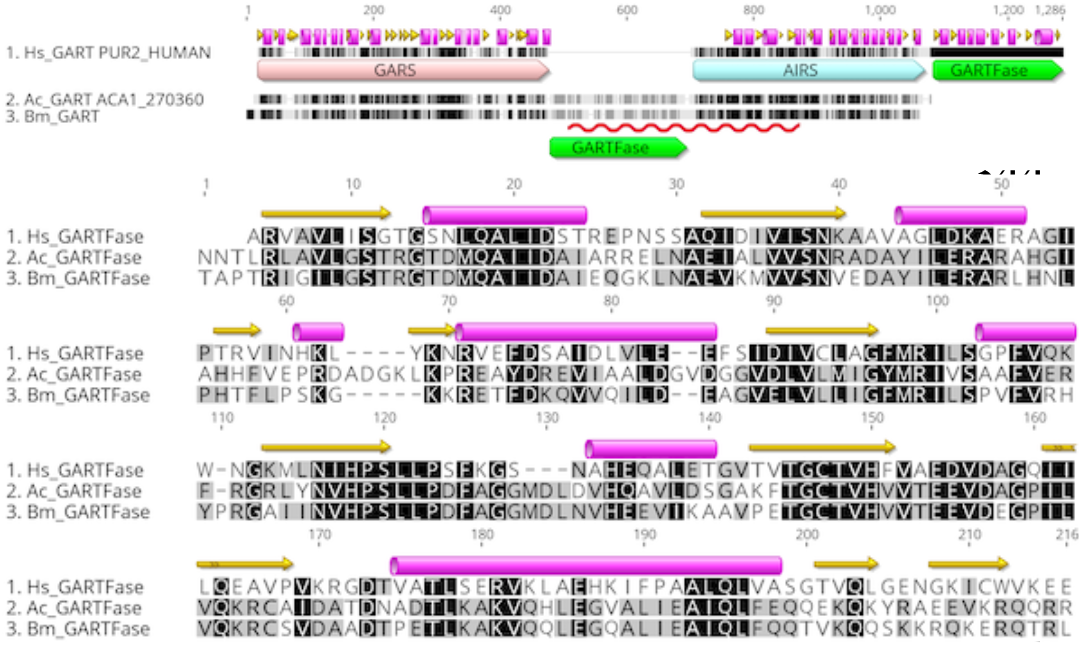
<u>Top:</u> Domain arrangement in human, *A. castellanii* and *B. mandrillaris* GART. Domains are annotated as large arrows on the alignment and higher level of residue conservation is represented as darker shades of gray. The region validated by direct sequencing in *Balamuthia* is underlined with a red squiggle. Secondary structure elements from human GART crystal structures are taken from UniProt. <u>Bottom:</u> Alignment of the GARTFase domains extracted from the GART sequences above.

## Conclusion

Through drug susceptibility screening with known antiparasitic compounds against *B. mandrillaris*, we identified protein targets with potential for treating *Balamuthia* granulomatous amoebic encephalitis. The reconstruction of the proteome from RNA-seq and annotation of the proteome allowed us to amplify, clone and validate the *B. mandrillaris* targets by direct sequencing. Our results indicate that the haploid proteome, consisting of the EviGene “main” proteins, is of high quality and provides an essential resource for further drug discovery and biological investigation. This study illustrates how the combination of phenotypic drug screening and a single RNA-seq experiment with short reads are enabling structure-based drug design against a eukaryotic pathogen with no prior proteome information.

## Materials and methods

### Cell culture

#### Maintenance of *Balamuthia mandrillaris*

The pathogenic *B*. *mandrillari*s (CDC:V039; ATCC 50209), a GAE isolate, isolated from a pregnant Baboon at the San Diego Zoo in 1986 was donated by Luis Fernando Lares-Jiménez ITSON University, Mexico^28^. Trophozoites were routinely grown axenically in BMI media at 37°C, 5% CO_2_ in vented 75 cm^2^ tissue culture flasks (Olympus), until the cells were 80-90% confluent. For sub-culturing, 0.25% Trypsin-EDTA (Gibco) cell detachment reagent was used to detach the cells from the culture flasks. The cells were collected by centrifugation at 4,000 rpm at 4°C. Complete BMI media is produced by the addition of 10 % fetal bovine serum and 125 μg of penicillin/streptomycin antibiotics. All experiments were performed using logarithmic phase trophozoites.

### Target identification

#### Phenotypic Screening

We previously developed and standardized robust high-throughput screening methods for the discovery of active compounds against *B. mandrillaris* trophozoites^28^. The trophocidal activity of compounds were assessed using the CellTiter-Glo 2.0 luminescent cell viability assay (Promega, Madison, WI). In brief, *B. mandrillaris* trophozoites cultured in BMI-complete media were seeded at 16,000 cells/well into 96-well plates (Thermo Fisher 136102) with various compounds diluted in 2-fold serial dilutions to determine the 50% inhibitory concentration (IC_50_). The highest percentage of DMSO diluted in the highest screening drug concentration was 1%. Control wells were supplemented with 1 % DMSO or 12.5 μM of chlorhexidine, as negative and positive controls, respectively. All assays were incubated at 37°C for 72 hours. At the end time point, 25 μL of CellTiter-Glo reagent was added to all wells. The plates were shaken using an orbital shaker at 300 rpm at room temperature for 2 minutes to induce cell lysis. After shaking, the plates were equilibrated at room temperature for 10 minutes to stabilize the luminescent signal. The ATP luminescent signal (relative light units; RLUs) were measured at 490 nm by using a SpectraMax i3X (Molecular Devices, Sunnyvale, CA). Drug inhibitory concentration (IC_50_) curves were generated using total ATP RLUs where controls were calculated as the average of replicates using the Levenberg-Marquardt algorithm, using DMSO as the normalization control, as defined in CDD Vault (Burlingame, CA, USA). Values reported are from a minimum of two biological replicates with standard error of the mean.

#### Selection of Target Genes

The protein names for verified potential targets were retrieved through Calibr at Scripps Research (https://reframedb.org/). The corresponding human protein sequences were downloaded from UniProt and queried against the *B. mandrillaris* assemblies using BLAST sequence similarity searches. Candidate targets were confirmed by comparing their protein sequences with closest sequence homologues in *Acanthamoeba* and *Naegleria* species and checking the *B. mandrillaris* functional annotation, where available. ORFs were selected for cloning from the Trinity *de-novo* assembly. Manual correction of putative start sites from multiple sequence alignments was performed with Geneious Prime 2019.1.1 (https://www.geneious.com).

### RNA extraction, library preparation and sequencing

#### RNA Extraction

*B. mandrillaris* were cultured and harvested as described above; the cells were counted and adjusted to 2 million cells for each extraction. Total RNA isolated using the RNA extraction kit (Agilent) as per manufacturing instructions. In brief, to the pellet of *Balamuthia* cells, 350 μl of lysis buffer and 2.5 μL of β-mercaptoethanol were added and homogenized. This was transferred into a prefilter spin cup and centrifuged at maximum speed, 14,000 x g, for 5 minutes. The filtrate was retained and an equal volume of 70% ethanol was added to the filtrate and vortexed until the filtrate and ethanol were mixed thoroughly. This mixture was then transferred into an RNA binding spin cup and receptacle tube and centrifuged at maximum speed for 1 minute. The filtrate was discarded and 600 μL of 1x low salt buffer was added and centrifuged at maximum speed for 1 minute. The filtrate was removed and centrifuged at maximum speed for 2 minutes. DNase solution was added and incubated for 15 minutes at 37°C. After incubation 600 μL of 1x high salt buffer (contains guanidine thiocyanate) was added and centrifuged at maximum speed for 1 minute. The filtrate was discarded and 300 μL of 1x low salt buffer was added and centrifuged at maximum speed for 2 minutes. 100 μL of elution buffer was added and incubated at room temperature for 2 minutes. Final elution was into a sterile 1.5 mL microcentrifuge tube at maximum speed for 1 minute.

Extracted RNA was stored at −80°C until further required. The integrity and purity of the RNA was assessed via RT-PCR and gel electrophoresis on a 2% agarose gel. The concentration was determined by measuring 280nM absorbance on a nanodrop (Nanodrop 1000, Thermo Scientific).

RNA quality was reassessed after a freeze thaw cycle using the Bioanalyzer RNA 6000 pico chip (Agilent, 5067-1513) and quantity was assessed using the Qubit RNA Broad Range Assay (Invitrogen, Q10210). The mRNA was isolated using the NEB Poly(A) mRNA Magnetic Isolation Module (NEB, E7490S) and prepared using a version of the Stranded RNA-seq protocol that was modified for *Leishmania*^40,41^. Only the negative stranded RNA-seq library preparation portion was performed. Library quantity and quality was assessed using the Qubit dsDNA High Sensitivity Assay (Invitrogen, Q32851), Bioanalyzer High Sensitivity DNA Chip (Agilent, 5067-4627) and the KAPA library quantification kit (Roche, KK4824). Libraries were sequenced on the Illumina Hiseq 4000, yielding 2 × 75 bp paired end reads.

### Transcripts assembly and annotation

Reads were quality filtered with Trimmomatic and assembled *de-novo* with Trinity v2.8 (k-mer=25) and Spades v3.13 (k-mer=29 and 33) after clipping of the adaptor sequences^42–44^. Further, quality-filtered reads were aligned to the published *B. mandrillaris* genome LFUI01 with STAR v2.6 and assembled with Trinity^45^. The three assemblies thus obtained were combined with EvidentialGene v19jan01 (EviGene) with BUSCO homology scores as input for the classifier^31^. Throughout the analysis, BUSCO v3 analysis was performed on either the European or Australian Galaxy mirrors ^46,47^. The Trinity *de-novo* assembly was functionally annotated with Trinotate^48^. Trinotate annotation sources included BLASTX and BLASTP homology searches against Swiss-Prot and AmoebaDB *A. castellanii*, PFAM domain analysis, as well as secretion and trans-membrane domain predictions with SignalP and TmHMM, respectively. Functional descriptions and gene ontology (GO) annotations of the EviGene ‘main’ proteins were predicted with PANNZER2^49^. GO annotations that were highest ranked by PANNZER2 were visualized with WEGO 2.0^50^.

### Comparison to other species and phylogenetic analysis

Proteome comparisons to other species in the UniProt database were obtained from the AAI-profiler server^51^. Cluster analysis and Venn diagrams of orthologous clusters were generated with OrthoVenn2^52^. Unless otherwise specified, all BLAST searches were performed with BLAST+ v2.8.1 and an expectation value of 0.001^53^. Pairwise distances for alignment-free phylogeny reconstruction were calculated with Prot-SpaM^54^. Input sequences included the *Balamuthia* EviGene ‘main’ proteins (this study), AmoebaDB *A.castellanii* strain Neff, *N. fowleri* ATCC 30894 Braker1 predicted proteins^55^ and UniProt complete reference proteomes (*C. fasciculata* UP000007797, *N. gruberi* UP000006671, *P. pallidum* UP000001396, *D. discoideum* UP000002195, *P. fungivorum* UP000241769 and *T. lacteum* UP000076078). Phylogenetic relationships were inferred by constructing a neighbor-joining tree from the word match-based Prot-SpaM distance matrix using MEGA X^56,57^.

### PCR and sequence validation

#### Cloning

All *B. mandrillaris* constructs were cloned, expressed, and purified using SSGCID established protocols^58,59^. The genes selected were PCR-amplified using cDNA template and purchased primers (Integrated DNA Technologies, Inc., Coralville, IA) (**Table S1**). The amplicons were extracted, purified and cloned into a ligation-independent cloning pET-14b derived, N-terminal His tag expression vector pBG1861 with a T7 promoter^60^. The cloned inserts were then transformed into purchased GC-5 cells (Genesee Scientific, El Cajon, CA) for ORF incorporation. Plasmid DNA was purified from the subsequent colonies and further transformed in chemically competent *E. coli* BL-21(DE3)R3 Rosetta cells with a chloramphenicol restriction.

#### Sequence Validation

Each *B. mandrillaris* construct was sequenced from both 5’- and 3’-ends with a custom forward primer (5’-GCGTCCGGCGTAGAGGATC-3’, 40nt upstream from the T7 promoter customary forward primer) and the T7 terminator reverse primer (5’-GCTAGTTATTGCTCAGCGG-3’) at GeneWiz (South Plainfield, NJ). The reads were assembled and matched to the expected sequences with the phrap assembler and cross_match^61^. Translations of the longest ORF in all six frames of the consensus sequence (or the forward read if unassembled) were then aligned using MUSCLE^62^ with the SSGCID target protein sequence to determine the best translated protein sequence and its alignment, percent identity and percent coverage. Manual examination of the sequences and alignments was performed in Geneious.

## Data availability

Illumina raw reads have been deposited at the National Center for Biotechnology Information (NCBI) BioProject repository with the accession number SRR12006108 under project PRJNA638697. The annotated protein sequences from the EviGene ‘main’ assembly are available on NIH Figshare under DOI: https://doi.org/10.35092/yhjc.12478733.v1. All data that are associated with the drug susceptibility study are archived using the database from Collaborative Drug Discovery (CDD; http://www.collaborativedrug.com/). The CDD database accommodates both compound chemistry data and results from phenotypic or target activity, cytotoxicity screening, and computed properties. The CDD database is becoming an established standard for the sharing of data within this community and we are eager to facilitate the distribution of our results in a similar manner.

## Notes

### Author contributions

Conceptualization, I.Q.P., C.A.R., D.E.K., and P.J.M.; methodology, I.Q.P., C.A.R, J.C., J.M., S.S. and L.T.; validation, I.Q.P. and C.A.R; formal analysis, I.Q.P., C.A.R., R.E.N., V.S., J.C. and S.S.; resources, D.E.K., W.C.V.V., J.C.M., and P.J.M.; data curation, I.Q.P. and C.A.R; writing-original draft preparation, I.Q.P. and C.A.R.; writing-review and editing I.Q.P., C.A.R., R.E.N., V.S., S.S., L.K.B., J.C.M., D.E.K., W.C.V.V., and P.J.M.; visualization, I.Q.P. and C.A.R.; supervision, I.Q.P., C.A.R., D.E.K., W.C.V.V. J.C.M., and P.J.M.; funding acquisition, D.E.K., W.C.V.V., J.C.M. and P.J.M. All authors have read and agreed to the published version of the manuscript.

## Acknowledgements

We thank Drs. Luis Fernando Lares-Jiménez & Fernando Lares-Villa (Instituto Tecnológico de Sonora, Ciudad Obregón, Sonora, Mexico) for the pathogenic isolate of *B. mandrillaris* used in this study. The authors acknowledge the support of the Freiburg Galaxy Team led by Prof. Rolf Backofen, Bioinformatics, University of Freiburg, Germany funded by Collaborative Research Centre 992 Medical Epigenetics (DFG grant SFB 992/1 2012) and German Federal Ministry of Education and Research (BMBF grant 031 A538A de.NBI-RBC). Rooksana Noorai was supported by an Institutional Development Award (IDeA) from the National Institute of General Medical Sciences of the National Institutes of Health under grant number P20GM109094. This project has been funded in part with Federal funds from the National Institute of Allergy and Infectious Diseases, National Institutes of Health, Department of Health and Human Services, under Contract No.: HHSN272201700059C and the Georgia Research Alliance (GRA) also supported this work.

